# Sponges as bioindicators for microparticulate pollutants

**DOI:** 10.1101/2020.05.26.116012

**Authors:** Elsa B. Girard, Adrian Fuchs, Melanie Kaliwoda, Markus Lasut, Evelyn Ploetz, Wolfgang W. Schmahl, Gert Wörheide

## Abstract

Amongst other threats, the world’s oceans are faced with man-made pollution, including an increasing number of microparticulate pollutants. Sponges, aquatic filter-feeding animals, are able to incorporate fine foreign particles, and thus may be a potential bioindicator for microparticulate pollutants. To address this question, 15 coral reef demosponges sampled around Bangka Island (North Sulawesi, Indonesia) were analyzed for the nature of their foreign particle content using traditional histological methods, advanced light microscopy, and Raman spectroscopy. Sampled sponges accumulated and embedded the very fine sediment fraction (< 200 µm), absent in the surrounding sand, in the ectosome (outer epithelia) and spongin fibers (skeletal elements), which was confirmed by two-photon microscopy. A total of 34 different particle types were identified, of which degraded man-made products, i.e., polystyrene, cotton, titanium dioxide and blue-pigmented particles, were incorporated by eight specimens at concentrations between 91 to 612 particle/g dry sponge tissue. As sponges can weigh several hundreds of grams, we conservatively extrapolate that sponges can incorporate on average 10,000 microparticulate pollutants in their tissue. The uptake of particles, however, appears independent of the material, which suggests that the fluctuation in material ratios is due to the spatial variation of surrounding microparticles. Therefore, sponges have a strong potential to biomonitor microparticulate pollutants, such as microplastics and other degraded industrial products.

## INTRODUCTION

Microparticulate pollutants (later referred to as “micropollutants”) are a threat to inhabitants of the world’s oceans. Here, we define micropollutants as man-made substances, or products of their subsequent degradation, smaller than 5 mm in size. They are introduced into the environment and are potentially harmful to organisms, for instance as microplastics, textile fibers and particulate toxins leach from household and cosmetic products (Dris et al. 2016; Auta et al. 2017; Rochman 2018). Because traditional sieving techniques fail to assess the very fine particulate fraction (< 200 µm) adequately (Lindeque et al. 2020), the main question driving this research is whether potential bioindicators for such anthropogenic micropollutants can be identified among marine organisms.

Sponges (Phylum Porifera) are aquatic benthic animals, which are geographically widely spread (Bell 2008). They consume mainly dissolved organic carbon (DOC), prokaryotes and ultraphytoplankton (< 10 µm) by filtering fine particles from the ambient water (Yahel et al. 2006). They incorporate particles following two main paths: (1) phagocytosis by choanocytes and (2) endocytosis through the exopinacoderm (Willenz and van de Vyver 1982; Teragawa 1986a). First, choanocytes, i.e., cells that generate the water flow in the sponge body through the beating activity of their flagellum, filter the water and capture the nutritive particles via phagocytosis (Hammel and Nickel 2014). The ingested particles are thought to be directly transferred from choanocytes to archaeocytes, which might be responsible for the digestion and transportation of nutrients to different sites in the sponge body (Willenz and van de Vyver 1982). Second, exopinacocytes may incorporate particles as big as 2 mm diameter, which deposited on the outside of the animal on the ectosome (Cerrano et al. 2002). Such microparticles may subsequently be transported by ameboid mesohyl cells from the ectosome towards sites of skeletogenesis in non-spiculated demosponges (Teragawa 1986a). This process is also associated with the spicule transport within the mesohyl by collencytes in spiculated Demospongiae (Teragawa 1986a). Foreign microparticles provide sponges with strength and support their growth (Teragawa 1985). They may also serve for protection (Burns and Ilan 2003) and anchorage to the substrate (Cerrano et al. 2002); Teragawa 1986b). However, mechanisms behind particle incorporation, retention and rejection in sponges are not fully understood yet. Nonetheless, we hypothesize that the fluctuation in material ratios incorporated by sponges is due to the spatial variation of surrounding microparticles; therefore, sponges may incorporate man-made micropollutants if present in their immediate environment and be viable models for biomonitoring such.

To address this issue, we carried out a combination of field and laboratory studies. The sampling of sponges was conducted in Indonesia since it is known to be a hotspot for land-based pollution in the middle of the Coral Triangle (Eriksen et al. 2014). Collected sponges were then transported to the home laboratories to locate, identify and quantify foreign particles, including micropollutants, within sponge bodies. We used histological methods, such as (nonlinear) light microscopy, as well as Raman spectroscopy for five poriferan species from Bangka Island (North Sulawesi, Indonesia) to address the following three questions: in which structure(s) do particles accumulate?; what kind of particles do sponges incorporate (diversity)?; do sponges have the potential to monitor microparticulate pollutants? Findings from this study contribute to fill a knowledge gap on particle incorporation by sponges. Indeed, sampled sponges appeared to be efficient sediment traps as they accumulated the very fine sediment fraction (< 200 µm), absent in the surrounding sand, in the ectosome and spongin fibers. A total of 34 different particle types were identified, of which degraded man-made products, such as polystyrene, cotton, titanium oxides and blue-pigmented particles, were incorporated by eight specimens. The incorporation of particles, however, appears independent of the material, which suggests that the fluctuation in material ratios is due to the spatial variation of surrounding microparticles. Therefore, sponges have a strong potential to biomonitor microparticulate pollutants, such as microplastics and other degraded industrial products.

## MATERIAL AND METHODS

### Site of study and sample collection

The field work took place at Coral Eye Resort on the west coast of Bangka Island (Kabupaten Minahasa Utara, Perairan Likupang), Sulawesi Utara, between March 17th and April 12th 2019, to assess the plastic contamination in marine sponges (research permit holder: Elsa Girard; SIP no.: 97/E5/E5.4/SIP/2019). The sampling area spanned approximately 7 km^2^ and specimens were sampled at three different locations: Coral Eye house reef, South and North of the jetty, and the mangroves (Supplementary material Fig. S1, Tab. S1). Non-lethal samples (n = 101) were taken on a wide variety of marine sponge species and sizes, using a stainless-steel diving knife, while snorkeling (free-diving). All sponges were sampled between 1 and 5 m water depth. Samples were collected in Kautex™ PVC wide neck square containers (250 and 500 mL) filled with ambient seawater until further treatment at Coral Eye Resort laboratory. An *in situ* picture of each specimen was taken, showing the macro-morphological features of the species. In addition, one sand sample from Coral Eye Resort was collected for comparison in the intertidal zone near the jetty (later referred to as beach sand).

### Specimen selection

A superficial inspection of the collected samples was done at Coral Eye Resort laboratory in order to assess the presence of foreign particles in the sponge tissue, potentially containing microplastics. Two techniques were used. In a first step, the samples were enzymatically digested with 0.2 mL Proteinase-K (20 mg/mL) mixed to 2 mL Tris-HCl buffer (0.1 mL Tris-HCL 1M, 0.02 g SDS 1%, 1.88 mL ddH_2_O) in a water bath (50-60 °C) for 3 h; a drop of the resulting solution was transferred on a microscope slide for inspection. In a second step, each sample was scrutinized with binoculars to search for apparent foreign particles within the sponge tissue. Specimens were selected depending on their foreign particle content for further downstream analysis, i.e., molecular species identification, histological and Raman spectroscopy analyses. Once a specimen-type was selected, two other specimens of apparently the same species were also sampled to create triplicates, i.e., 3 specimens of the same species. A total of 15 specimens were sampled, i.e., from five different species, for further downstream analysis. Collected samples were preserved in two aliquots: 96% ethanol for DNA barcoding and 4% formaldehyde for histology and spectroscopy.

### Species identification

At the Molecular Geobiology and Paleobiology laboratory of the Department of Earth & Environmental Sciences, Paleontology & Geobiology, LMU Munich, specimens were identified to the genus using an integrative approach (Wörheide and Erpenbeck 2007; Voigt and Wörheide 2016). The DNA was extracted from the samples using a DNA extraction kit (NucleoSpin® Tissue, Macherey-Nagel GmbH & Co. KG). DNA barcoding was conducted using a fragment of the 28S ribosomal DNA, a region amplified using universal primers via polymerase chain reaction (PCR) (Supplementary material Tab. S2). PCR products were run through an agarose gel (1%) to verify that the amplification was correctly performed and amplified fragments were of expected length (ca. 550 bp). The DNA was sequenced with BigDyeTerminator v3.1. Sanger Sequencing was conducted at the Genomic Sequencing Unit of the LMU Munich, using an ABI 3730 (Erpenbeck et al. 2017).

Forward and reverse sequences were assembled and edited using CodonCode Aligner v3.7.1.2 (www.codoncode.com). Sequences of poriferan origins were identified with BLAST® for nucleotides using the NCBI database (https://blast.ncbi.nlm.nih.gov). Finalized sequences were combined to the 28S sponge data set (Erpenbeck et al. 2016) available at the Sponge Genetree Server (www.spongegenetrees.org). The data set was largely reduced to concentrate on the important clades, by selecting only the nearby taxa (taxonomically classified) with the least genetic distance to the samples. Alignments were performed in MAFFT v7.427 (https://mafft.cbrc.jp/alignment/software/), default settings. Subsequently, phylogenetic trees were calculated for 28S sequences in Seaview v4.6.3 (Gouy et al. 2010) under PhyML GTR+G+I model with the invariable site and gamma shape settings obtained via jmodeltest 2.1.10 v20160303 (Darriba et al. 2012), and included 100 bootstrap replicates (Guindon et al. 2010). FigTree v1.4.3 (http://tree.bio.ed.ac.uk/) and Adobe Illustrator CS3 were employed to modify the esthetic of the figures. Final barcoding data (alignments and trees) is stored on GitHub repository (https://github.com/PalMuc/PlasticsSponge).

### Histological analysis

The samples were fixed with 4% formaldehyde and gradually dehydrated with ethanol at the Coral Eye Resort laboratory (Indonesia). At the molecular laboratory in Munich, samples were prepared for thin sectioning in LR-white medium to preserve the original position of foreign particles within the tissue. Sections with a thickness ranging between 50 to 400 µm were cut depending on the specimen morphology, using a saw microtome (Leica SP1600). Two freshly cut sections per sample (30 to 50 µm thick) were stained using Carbol Fuchsin dye (Roth staining kit) to highlight the cellular distribution within the mesohyl, including choanocyte chambers. Sections were mounted on microscope slides using Eukitt Quick-hardening mounting medium. The histological analysis was conducted using a microscope Leica DMLB (Type 020-519.502 LB30 T BZ:00, Leica Mikroskopie & Systeme GmbH Wetzlar) with a mounted digital camera. Images of the same field of views were taken under brightfield and (cross-)polarized light illumination. The polarized light fields allowed a better recognition of the foreign particles in the sponge, embedded in the organic tissue. Field of views of the ectosome, mesohyl, skeletal structures and aquiferous system (i.e., canals and choanocyte chambers) were recorded for each specimen. The histological analysis also enabled the description of the sponge main morphological micro-features.

In addition to the assessment of particle accumulation areas, relative particle abundance and size were analyzed with ImageJ v1.52K (Schindelin et al. 2012). All tissue images utilized for the analysis were taken under the same settings (100 µm thin section, equal luminosity and magnification) to ensure comparability of the data. To measure the particle’s relative abundance, images were translated into grayscale (8-bit) and the mean light intensity of ten random square areas (264.52 µm side length) were measured per structure and per sample. The mean light intensity is a numerical value generated by the software that allows for a comparison between areas and samples with an arbitrary unit (AU). Unpolarized light illumination was chosen for keratose sponges, because the spongin tissue from the skeletal fibers has a high transmission with little scattering comparable to the particles. On the contrary, (cross)polarized light illumination was used for heteroscleromorphs, to observe particles in these heavily spiculated specimens with high organic matter content. Images taken with polarized light were treated in a second step by inverting the gray scale in order to have dark particles on a pale background, similarly to the images taken with unpolarized light. Therefore, the lower the intensity value, the darker the area and the more particles are present. The particle size was also assessed and categorized with ImageJ: small (most particles <50 µm diameter), large (most particles > 50 µm diameter) or mix (presence of small and large particles at a similar fraction). The data gathered from the particle distribution, abundance and size was analyzed in R v3.3.3 (R Core Team 2017). Primary data and R scripts are available on GitHub repository (https://github.com/PalMuc/PlasticsSponge).

### Particle distribution with two-photon excitation

Histological sections of sponge tissue were analyzed by two-photon excitation (TPE) 3D imaging, to highlight the contrast between the highly fluorescent organic tissue from the sponges in opposition to the non-fluorescent mineral particles incorporated by the sponge. Samples were evaluated after histological preparation. For two-photon imaging, fresh sections of 170 µm thickness were prepared without staining and mounted on microscope slides using Eukitt Quick-hardening mounting medium. Brightfield pictures of the scanned area were taken before the experiment followed by a 3D scan of the specimen. Each 2D image had a range of 190 µm, an acquisition time of 180 s and a step size of 380 nm. The 3D step size between the 2D image planes and the total number of planes was chosen with respect to the object of interest and ranged from 10 to 21 planes with 0.5 to 3 µm steps.

Imaging was carried out on a confocal scanning microscope (TE 300; Nikon) with mounted bright-field illumination and camera. The two-photon excitation source was a fiber-based, frequency-double erbium laser (FemtoFiber dichro bioMP, Toptica Photonics) running at 774 nm. The laser power was 10 mW. The laser light was coupled into the microscope via a low pass dichroic mirror (HC BS 749 SP; AHF Analysetechnik) that separates laser excitation and fluorescence emission. Scanning of the sample in 3D was achieved by using a xyz piezo stage (BIO3.200; PiezoConcept). The laser excitation was focused onto the sample with a 60x (water) 1.20-NA plan apochromat objective (Plan APO VC 60x 1.2 NA, Nikon). The emission was collected by the same objective and passed afterwards through a bandpass filter (SP600; AHF). The emission was recorded on an APD detector (Count Blue; Laser Components) and its photons stream registered using a TCSPC card (TH260 pico dual; PicoQuant GmbH). The experiment was controlled using a home-written program written in C#. The confocal data was extracted and evaluated afterwards by PAM (Schrimpf et al. 2018) and ImageJ2 (Schindelin et al. 2012).

### Raman spectroscopy

Raman spectroscopy method permitted the identification and quantification of particles on a filter and *in situ*, i.e., from thin sections. As preparation for Raman measurements, samples were firstly subsampled, dried and weighed. The subsamples weighed between 2.2 and 11 mg. The sponge tissue was digested in 1.5 mL household bleach over 2-3 days, with a one-time bleach renewal. Subsamples were then washed with MilliQ water five times in a row. Particles left were filtered through a nitrocellulose membrane (Whatman™, 1 µm mesh size) with the aid of a vacuum pump. One hundred particles were randomly measured per sample (referred to as “random” search pattern). Furthermore, a maximum of ten additional particles per sample were measured by purpose depending on differences in color, shape and texture to assess the diversity of incorporated particles in lower abundance (referred to as “target” search pattern). The same method was performed on the beach sand sample. In addition, the spectrum of white and a red sclerites of *Tubipora musica* (sample number GW1858, obtained from an individual grown in aquarium at the Molecular Geobiology and Paleobiology laboratory) was measured to compare its red pigment signal to that of the red particles present in sponges. Moreover, one thin section (30-50 µm thick) per triplicate was chosen to confirm the presence of *in situ* particles, which are not organic matter. Five to six particles were measured in each structure for all five sections.

Raman spectra were taken on a confocal Raman microscope (HORIBA JOBIN YVON XploRa ONE micro Raman spectrometer) belonging to the Mineralogical State Collection Munich (SNSB). The used Raman spectrometer is equipped with a Raman edge longpass filter, a Peltier cooled CCD detector and three different lasers working at 532 nm (green), 638 nm (red) and 785 nm (near IR). Here, 532 nm excitation was used to perform the measurements, with a long working distance (LWD) objective magnification 100x (Olympus, series LMPlanFL N), resulting in a 0.9 µm laser spot size on the sample surface. The set power required for high-quality spectra varied between 10% and 100% (i.e., respectively 0.879 mW and 8.73 mW +/- 0.1 on the sample surface) depending on the type and size of measured particles. The diameter of pin-hole and the slit width were set to 300 and 100 µm, respectively. Each acquisition included two accumulations with an integration time of 8 s over a spectral range of 50 to 2000 cm^-1^ (ca. 35 s per measurement). Resulting Raman spectra were analyzed using LabSpec Spectroscopy Suite software v5.93.20, compiled in a table, visualized in R v3.3.3, manually sorted in Adobe Illustrator CS3, and compared with available spectra from RRUFF database (see: http://rruff.info/index.php) and published work (e.g., (Zięba-Palus and Michalska 2014). The statistical ANOSIM test (Analysis of Similarity) was performed in R v3.3.3, using 999 permutations in the vegan package (Oksanen et al. 2017) to assess the similarity in foreign particle assemblage composition between species, subclasses and sampling locations. Raman spectra and analysis scripts written in R used for the analysis are available on GitHub repository (https://github.com/PalMuc/PlasticsSponge).

### Precautions against contaminants

To avoid contamination of the samples, latex gloves, glassware, cotton towel and dust-free wipers (Kimtech Science) were used when manipulating the samples at all times. All open manipulations done in Molecular Geobiology and Paleobiology laboratory at LMU, i.e., dissection and filtration of the samples, were conducted under a clean bench (BDK, Luft- und Reinraumtechnik GmbH). Eppendorf safe-lock tubes 1.5 mL (polypropylene) were used to centrifuge the subsamples during tissue digestion and subsequent washing steps. Consequently, a negative sample was included, undergoing the same steps as all sponge samples from tissue digestion to particle filtration. Due to the airborne exposure of the samples in the field at Coral Eye Resort laboratory and the presence of fibers on the negative filter, all resulting fibers in this study were regarded as contaminants and therefore not taken into account. Only microparticles, excluding microfibers, were analyzed.

## RESULTS

Fourteen out of 82 morphologically different specimens (17%) contained foreign particles based on the superficial analysis conducted at Coral Eye Resort laboratory. Five particle-bearing species were particularly abundant on site (Bangka Island, Indonesia), and sampled three times to generate the triplicates (n = 15). In order to determine the accumulation areas of foreign particles in coral reef sponges, the 15 samples were taxonomically classified to the genus level into 5 clades: *Carteriospongia, Ircinia* I, *Ircinia* II, Tethyid I and Tethyid II (Supplementary material Fig. S2, Tab. S3) and histologically analyzed. Subsequently, the samples were examined with Raman spectroscopy to assess the diversity of incorporated particles.

### I. Particle distribution

Thin sections provided an overview of the main structures and the distribution of particles within sponge bodies. Incorporated particles were located and identified with polarized light microscopy using TPE 3D imaging and Raman spectroscopy. Choanocyte chambers were difficult to distinguish from the mesohyl and canals despite the staining. Moreover, no particles were found inside the visible choanocyte chambers; only particles surrounding choanocyte chambers were observed in both Tethyid species. Therefore, no further statistical analysis was conducted on choanocyte chambers.

Foreign particles were observed by (nonlinear) light microscopy in the aquiferous system, the mesohyl, the ectosome and the fibrous skeleton, depending on the species (Fig. 1, 2). Two-photon microscopy clearly confirmed that the incorporated particles were completely embedded in the surrounding tissue (Fig. 2, Supplementary material Fig. S3). All specimens incorporated particles in the mesohyl and the ectosome. Moreover, 73% of the specimens, independently of the skeletal material, had some particles in the aquiferous system; the clades *Ircinia* I did not contain particles in their canals. All specimens of the subclass Keratosa incorporated particles in their skeletal spongin fibers (e.g., Fig. 2D), whereas heteroscleromorphs did not in relation to their spiculated skeleton (Fig. 1).

**Figure 1.**
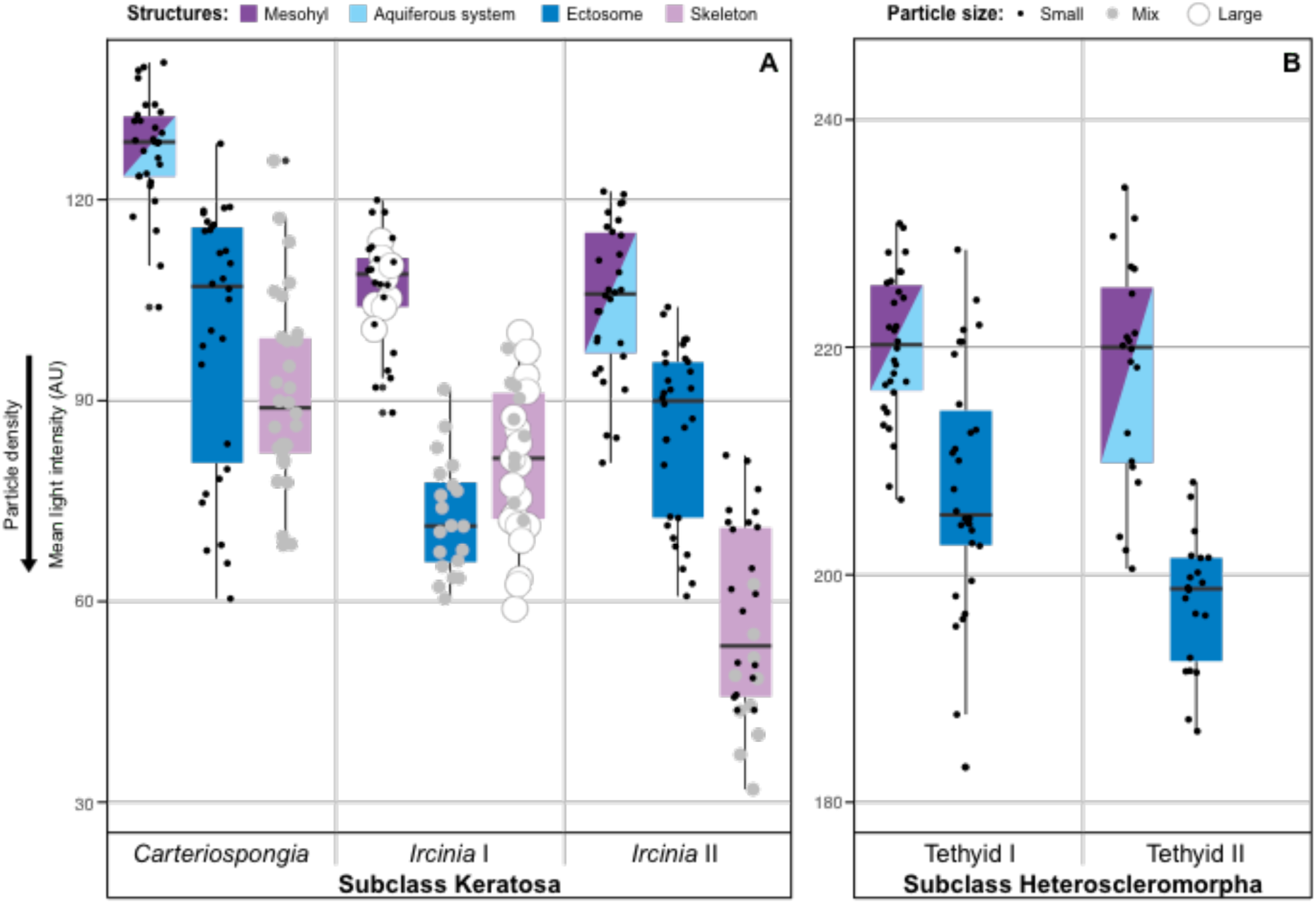
Particle distribution amongst sampled sponges and their abundance in tissue structures as derived from brightfield imaging. A) Subclass Keratosa, B) subclass Heteroscleromorpha. The lower the intensity, i.e., absorption of the specimen body, the higher the number of particles in the structure; the aquiferous system was not differentiated from the mesohyl in cases where particles were present in the aquiferous system. Particle size is represented by the size and color of the dots (i.e., black small dots for a majority of particles < 50 μm, white large dots for a majority of particles > 50 μm and gray medium dots for a more or less equal presence of small and large particles). Color code: skeleton in lila, aquiferous system in cyan, mesohyl in purple, ectosome in blue. The black horizontal line inside the boxes represents the median intensity value of the data. The mean light intensity (y-axis) is given in an arbitrary unit (AU). Note: relative intensity within the same subclass can be compared, however not between the subclasses as the scale used was different.

**Figure 2.**
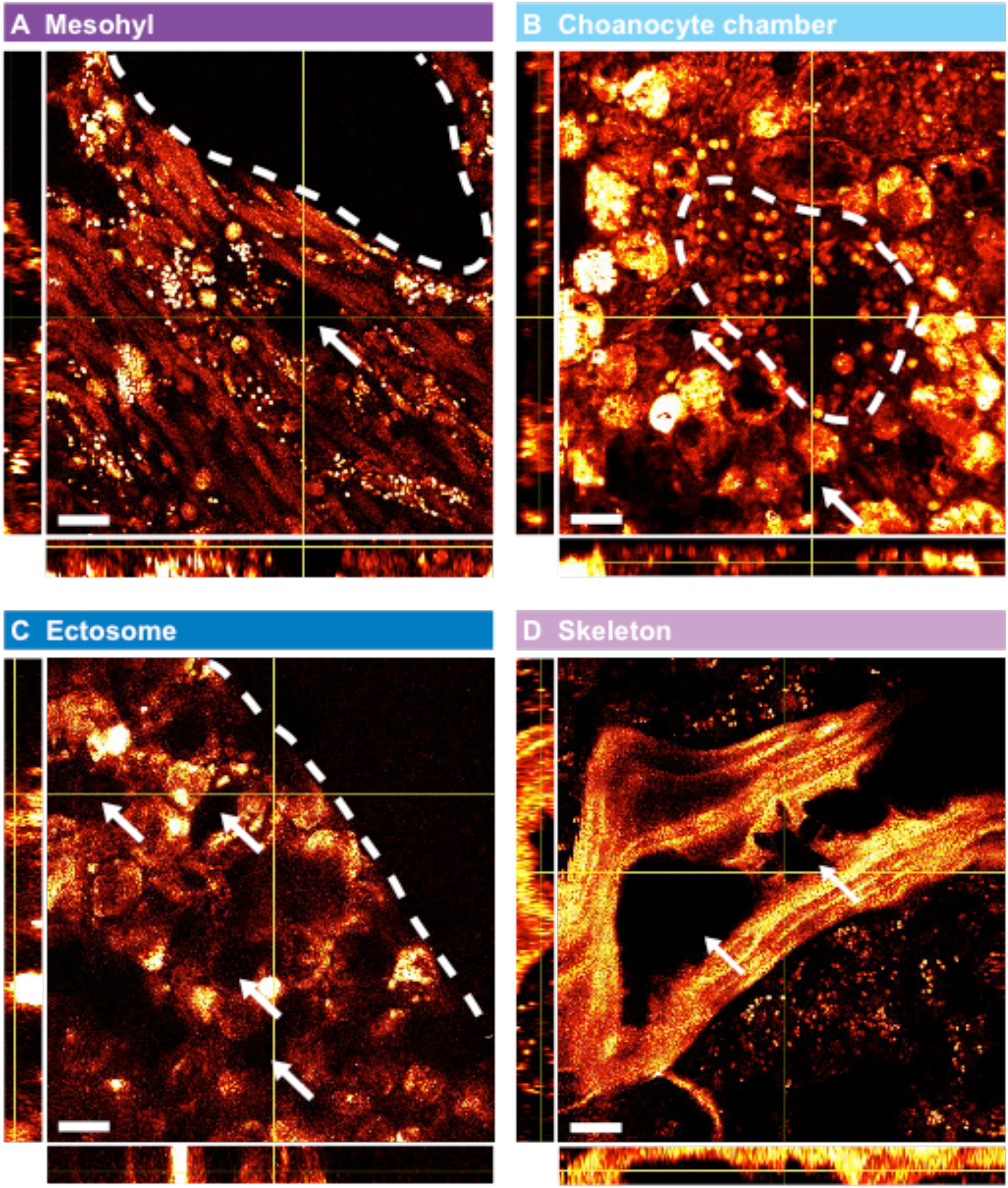
Two-photon images of three-dimensional sponge tissue sections with embedded foreign particles. The auto-fluorescence of the organic tissue material serves as imaging contrast compared to inorganic particles that are non-fluorescent. XZ and YZ projections through the 3D image stack are depicted aside of each XY projection (brightfield images are shown in Supplementary material Fig. S3). Exogenous particles are embedded in the tissue (marked with white arrows) and are found A) in the mesohyl of *Ircinia* sp. (the canal is circled with a dashed line), B) surrounding the choanocyte chamber of Tethyid sp. (circled with a dashed line), C) at the ectosome of Tethyid sp. (the dashed line separates the outermost part of the sponge tissue), and D) in spongin fibers of *Carteriospongia* sp.. Z-scan ranges are 18 µm and scale bars are 20 µm.

Across all specimens, particles embedded in the mesohyl (e.g., Fig. 2A) were in lower abundance (the higher the light intensity, less dense the particle cover), in comparison to the density of particles accumulated in spongin fibers and/or the ectosome (Fig. 1). Moreover, thorough microscopy analyses reveal a majority of small particles (< 50 µm) present in the mesohyl/aquiferous system and accumulated in the ectosome (Fig. 1, 2). Larger particles (> 50 µm) were observed in spongin primary fibers of keratose sponges (Fig. 1, 2). Only specimens from the clade *Ircinia* I incorporated large particles in all three structures. The size of uptaken particles between *Ircinia* species differed although they belong to the same genus. In fact, the clade *Ircinia* II reflected *Carteriospongia*’s particle size pattern. Measured particles on the filters varied in size, with a diameter ranging from 5 µm to approximately 200 µm, which are equivalent in size to the particles measured *in situ* (Fig. 2). The size of the particle may therefore indicate in which structure particles might have been incorporated in relation to Fig. 1, for example, larger particles in Keratosa are more likely to come from the fiber network than the mesohyl (Fig. 1, 2). The fine fraction (< 200 µm) was absent from the beach sand sample, where only coarse particles (> 500 µm diameter) were observed.

### II. Particle diversity

A total of 1,686 particles were measured on 15 filters (between 103 and 110 particles per filter). Across all measured particles on the filters, 34 different spectra were identified, of which 22 were associated to a single material or pigment (aragonite, calcite, amorphous calcite, quartz, ß-quartz, anatase, feldspar, graphite, magnetite, mackinawite, ferrosilite, cotunnite, hematite, riebeckite derivative, polystyrene, cotton, carbon, *Argopecten irradians* shell, red biomineralization type I and II, blue pigment type I and II) (Fig. 3, Supplementary material Tab. S4). The 12 remaining spectra are polymineralic particles and were interpreted as a mixture of two different materials (e.g., aragonite + quartz). Tab. 1 illustrates the variability and diversity of the scarce incorporated compounds between the different species.

**Table 1.** Number of scarce inorganic, organic and pigmented particles, which have been identified with Raman spectroscopy, including polyminerals but excluding calcitic or aragonitic particles. Numbers of articles and dry weight were summed between the specimens of the same species. Total dry weight of digested sponge tissue and the number of potentially anthropogenic particles are highlighted in gray.

|  |  | Number of particles found per species |  |  |  |  |  |  |  |  |  |
| --- | --- | --- | --- | --- | --- | --- | --- | --- | --- | --- | --- |
|  |  | *Carteriospongia* |  | *Ircinia* I |  | *Ircinia* II |  | *Tethyid* I |  | *Tethyid* II |  |
| Total dry weight of tissue |  | 12.5 mg |  | 18.7 mg |  | 14.8 mg |  | 28.1 mg |  | 18.6 mg |  |
| Search pattern |  | Random | Target | Random | Target | Random | Target | Random | Target | Random | Target |
| Inorganic | Quartz | 2 | --- | 3 | --- | 3 | --- | 8 | --- | 2 | --- |
|  | β-Quartz | --- | --- | --- | --- | --- | --- | --- | 1 | --- | 2 |
|  | Feldspar | 1 | --- | --- | --- | --- | 1 | 3 | --- | 3 | --- |
|  | Carbonate phosphate | --- | --- | --- | --- | 4 | --- | --- | --- | --- | --- |
|  | Anatase | --- | --- | --- | --- | --- | 1 | --- | --- | 2 | --- |
|  | Riebecktite derivative | --- | --- | --- | --- | --- | --- | --- | --- | 1 | --- |
|  | Magnetite | --- | --- | --- | --- | --- | --- | --- | --- | 1 | --- |
|  | Ferrosilite | --- | --- | 2 | 1 | --- | --- | 1 | --- | --- | --- |
|  | Hematite | --- | --- | --- | --- | --- | 1 | 1 | 1 | --- | --- |
|  | Cotunnite | 1 | --- | --- | --- | --- | --- | --- | --- | --- | 1 |
|  | Graphite | --- | --- | --- | --- | --- | --- | --- | --- | --- | 1 |
|  | Mackinawite | --- | --- | --- | --- | --- | --- | --- | 1 | --- | 2 |
|  | Quartz + Aragonite | 1 | --- | --- | --- | 1 | --- | 2 | --- | 1 | --- |
|  | Quartz + Anatase | --- | --- | --- | --- | 1 | --- | 3 | --- | 1 | 1 |
|  | Anatase + Feldspar | 1 | --- | 1 | --- | --- | --- | --- | --- | 1 | --- |
|  | Anatase + Calcite | --- | --- | --- | --- | --- | --- | 1 | --- | 1 | --- |
| Organic | Carbon | --- | --- | --- | --- | --- | --- | --- | 1 | --- | 2 |
|  | Cotton | --- | --- | --- | 2 | --- | 4 | --- | --- | --- | --- |
|  | Polystyrene | 1 | --- | 1 | --- | --- | --- | --- | --- | --- | --- |
| Pigment | Red pigment type III | --- | 3 | 1 | --- | --- | --- | --- | --- | --- | --- |
|  | Blue pigment type I | --- | --- | --- | --- | --- | --- | --- | 1 | --- | --- |
|  | Blue pigment type II | --- | --- | --- | --- | --- | --- | --- | 1 | --- | --- |

**Figure 3.**
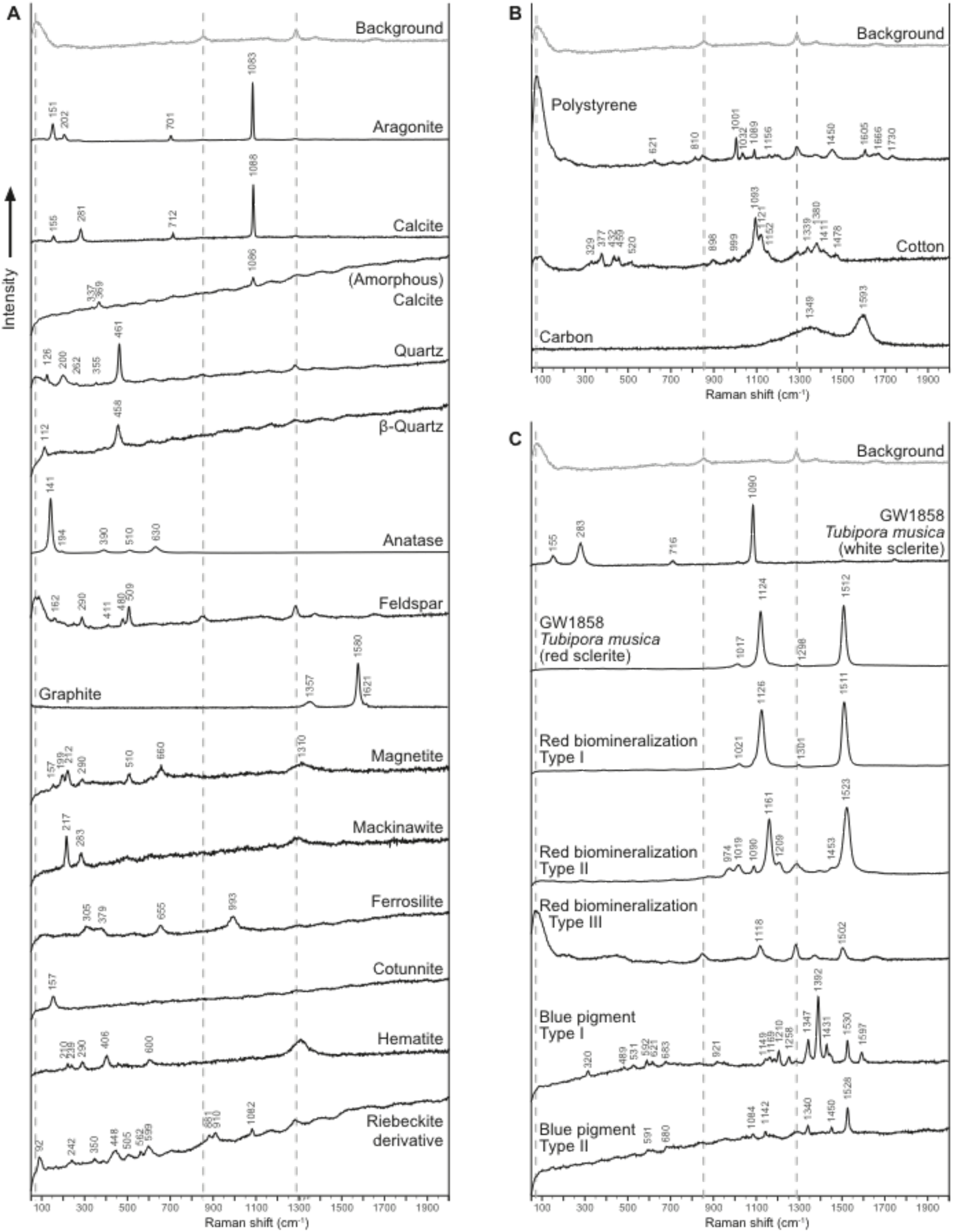
Raman spectra of incorporated particles (n = 1686). Mix spectra are not included, as they are a combination of two different minerals illustrated above. A) inorganic spectra. B) organic spectra. C) spectra associated with pigments. Red biomineralization spectra are compared to GW1858 (octocoral *Tubipora musica* red and white sclerites). Numbers over the signatures indicate the highest point of each vibrational band. Dotted lines indicate bands corresponding to the background.

#### Inorganic compounds

Independent of their identical chemical formula (i.e., CaCO_3_), calcite was distinguishable from aragonite with Raman spectroscopy due to a different crystal, i.e., trigonal vs. orthorhombic, respectively. This crystallographic change is translated into different vibrational bands in the lattice modes. Calcite and aragonite showed similar vibrational bands around 150, 705 and 1085 cm^-1^, but calcite had a band at ca. 280 cm^-1^, whereas aragonite had one at ca. 205 cm^-1^. Quartz was characterized with a vibrational band at 460 cm^-1^, related to the Si-O-Si bond. Carbonate phosphate showed two weak vibrational bands at ca. 960 cm^-1^ (phosphate; PO_4_^3-^) and 1075 cm^-1^ (carbonate; CO_3_^2-^). The latter is very close to the strongest vibrational band observed for calcite and aragonite (ca. 1085 cm^-1^), also associated with CO_3_^2-^. Feldspar was distinguished by three vibrational bands: ca. 290, 480 and 510 cm^-1^ (silica; Si), and anatase by one main band at ca. 140 cm^-1^. The D-band (ca. 1350 cm^-1^) and G-band (ca. 1580 cm^-1^) are indicative for carbon-based materials, such as carbon, graphene and graphite. Given the band shape and ratio between both lines (Roscher et al. 2019), we identified the compounds as carbon and graphite (Fig. 3).

#### Organic compounds

Polystyrene was also measured for two particles during a random search pattern. Particles, such as cotton and blue colored particles, were found during a target search pattern (Tab. 1). Some of these materials potentially come from man-made products, which is discussed in detail below. One specimen of *Carteriospongia* sp. (4.6 mg dry weight) and one of *Ircinia* I (6.3 mg dry weight) contained polystyrene at a concentration of 0.217 and 0.159 particle/mg, respectively. Four specimens from the Genus *Iricnia* (4.8, 4.9, 6.3 and 6.5 mg dry weight, respectively) were found with cotton at concentrations between 0.154 and 0.612 particle/mg.

#### Pigmented compounds

The Raman signature of a white and a red sclerite of *Tubipora musica* (GW1858) showed that the red pigment signal generally (peaks at 1326 and 1511 cm^-1^) covers all vibrational bands of calcite (ca. 280 and 1085 cm^-1^). However, this red pigment (type I), found also in red aragonitic particles (e.g., red type I + aragonite), did not overtake the vibrational bands of aragonite (ca. 150 and 1085 cm^-1^), ubiquitous across all samples. A second and third red pigments were also detected with Raman spectroscopy, showing peaks at 1161 and 1523 cm^-1^ and at 1118 and 1502 cm^-1^, respectively. Red pigment type III is slightly shifted compared to the type I and II (Fig. 3). Finally, blue-pigmented particles were observed on the filter of two specimens from the clade Tethyid I (11.0 and 8.5 mg dry weight, respectively) at a concentration of 0.091 (type I) and 0.118 (type II) particle/mg.

#### Mineral ratios

Aragonitic and calcitic particles were identified in all observed structures. Tethyid clades did not only incorporate minerals, but also a considerable amount of particulate organic matter (Fig. 2C, D). Keratosa specimens incorporated on average 68% aragonite and 25% calcite, whereas Heteroscleromorpha specimens had 59% and 30%, respectively (Fig. 4). The aragonite-calcite median ratio of Tethyid I was 1.875, Tethyid II 2.462, *Cartegiospongia* 2.680, *Ircinia* I 2.310 and *Ircinia* II 2.783. Although all specimens had a higher aragonite-calcite ratio on average than that of the beach sand (ratio = 1.735), no species had significantly different particle ratio compared to one another and to the sand sample (ANOSIM: *P* = 0.067) as well as between the subclasses (ANOSIM: *P* = 0.392) based on the Analysis of Similarity (Fig. 5C, D). Specimens were also compared on their aragonite-calcite ratio according to the location they were collected, i.e., Coral Eye house reef South or north. The Analysis of Similarity also indicates that the particle ratio did not significantly vary between the sampling sites (ANOSIM: *P* = 0.620) (Fig. 5E).

**Figure 4.**
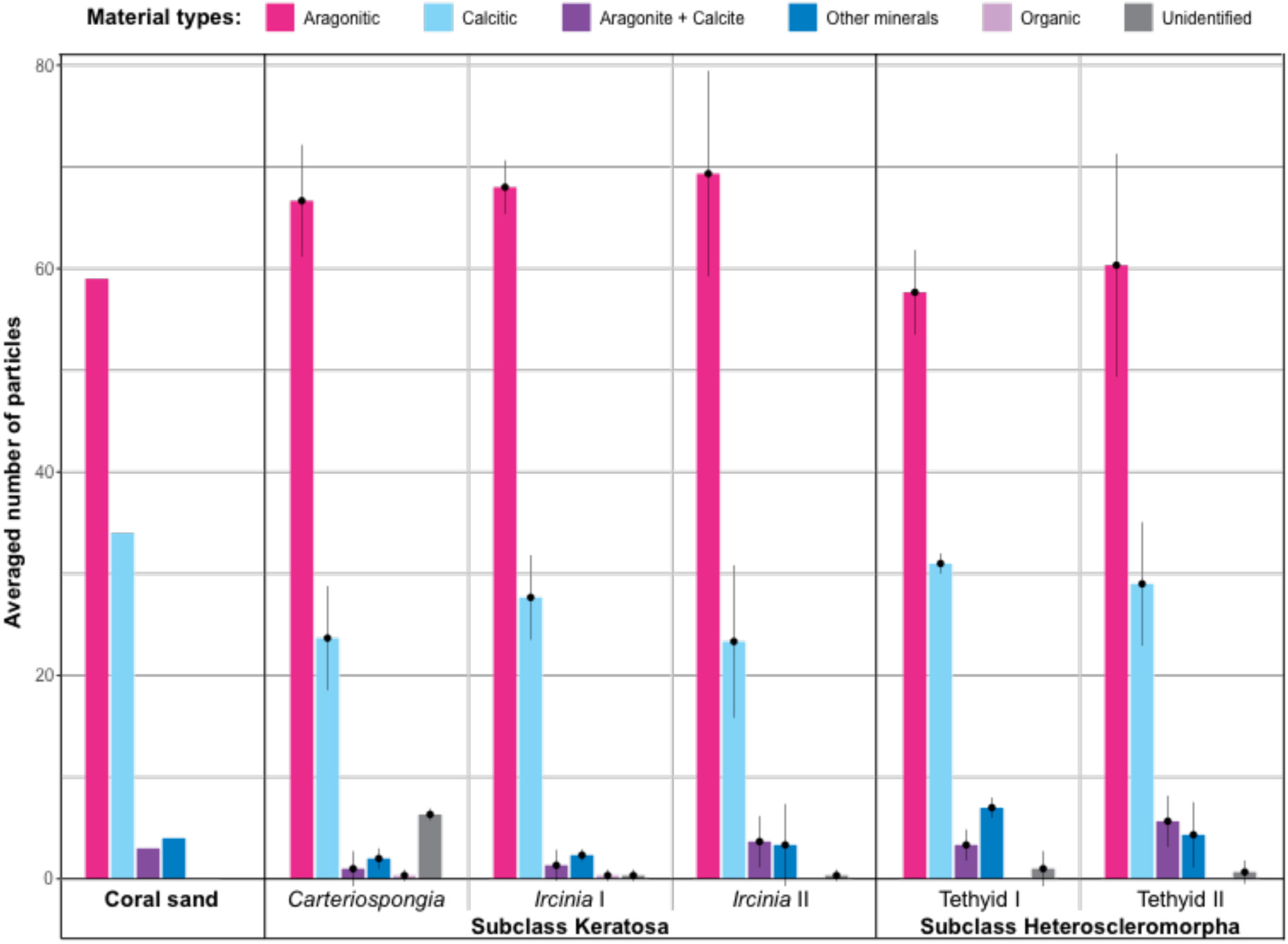
Diversity of materials among sampled sponges. Particle diversity from resulting Raman measurements of 100 randomly selected particles per sample. Each species accounts for 3 samples, and particle counts were averaged per clade (pink: aragonitic; cyan: calcitic; purple: aragonite + calcite; blue: other minerals; lila: organic; gray: unidentified). Error bars illustrate the variation in number of particles per category between the specimens of the same species. The category “Unidentified” results from low quality spectra, which could not be associated with any known material.

**Figure 5.**
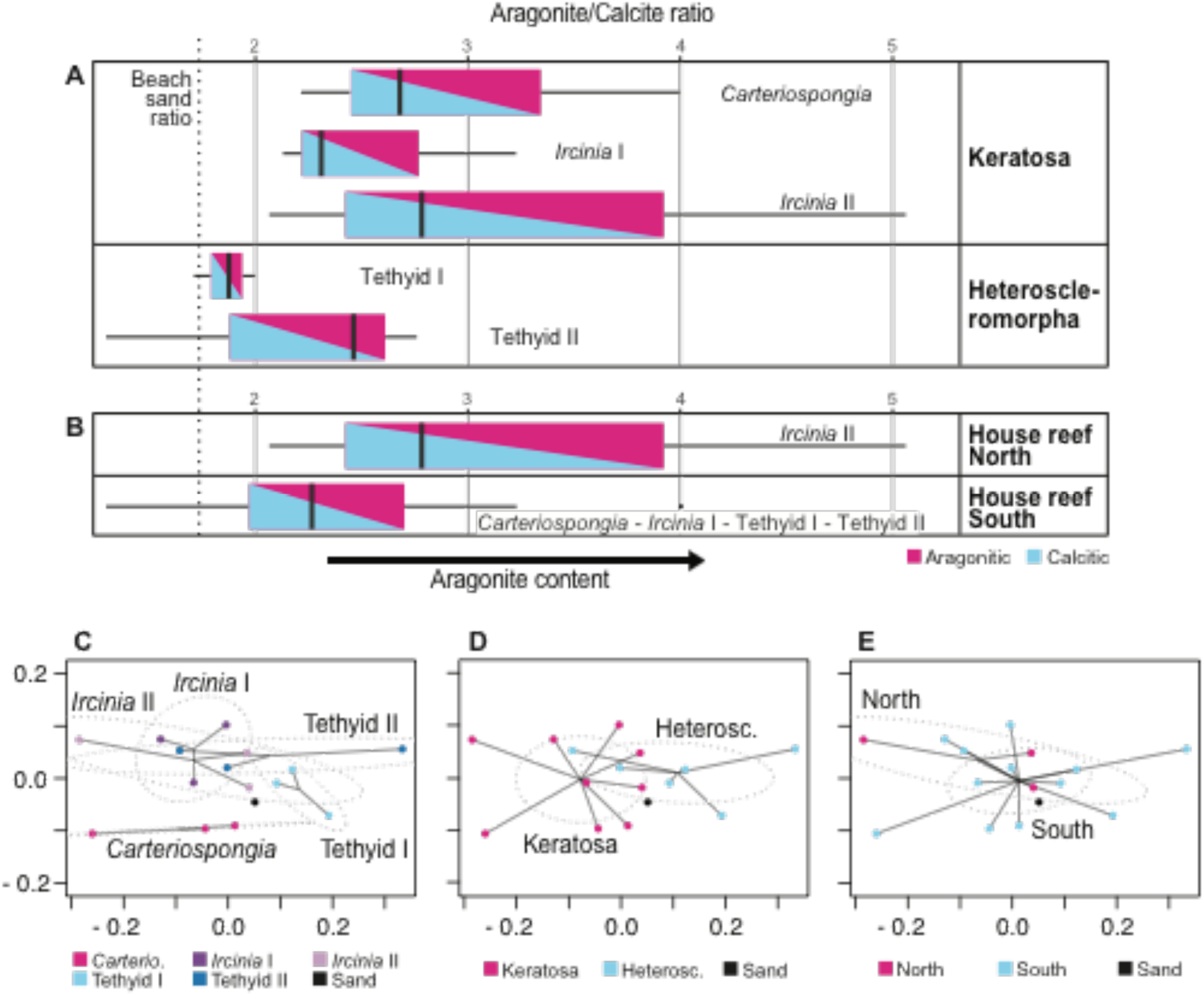
Aragonite to calcite ratio A) at the species level and B) according to the sampling site. The beach sand sample ratio (1.735) is indicated with the dotted vertical line. Non-metric multidimensional scaling (NMDS) plots C) at the species level, D) at the subclass level and E) according to the sampling site, displaying the differences between the foreign particle assemblage composition. Dashed gray ellipses represent the 99% confidence interval. Note: the Analysis of Similarity indicates no significant differences between the samples at all three comparison levels (ANOSIM: *P* > 0.05).

Other than aragonite and calcite, < 2% of quartz was measured across all clades; it was however not present in the beach sand. However, 4% of the coarse grains in the beach sand sample was feldspar, which was also identified in *Carteriospongia* and both Tethyid clades (Tab. 1). Carbonate phosphate and titanium oxide (anatase) were respectively measured on *Ircinia* II and Tethyid II filters (Tab. 1). Anatase was found at concentrations of 0.208 and 0.345 particle/mg. Most abundant polymineralic particles observed across the samples were quartz + aragonite and quartz + anatase, although both of them found in low concentrations (Tab. 1). The Analysis of Similarity suggests that the species have no preference on the material to be incorporated, because the species did not have significantly different material assemblage composition (ANOSIM: *P* > 0.05).

## DISCUSSION

In this study, 15 demosponges (*Carteriospongia* sp., *Ircinia* spp. and Tethyid spp.) were histologically analyzed and characterized with respect to their foreign particle content with light microscopy and Raman spectroscopy. The particle density was higher in the ectosome and spongin fibers of keratose sponges than in the mesohyl. No particles were observed in choanocyte chambers, but only surrounding them in Tethyid specimens. Embedded foreign particles were of larger size in keratose spongin fibers, whereas generally smaller than 50 µm in the mesohyl and the ectosome, as confirmed with TPE analysis. Keratose and heteroscleromorph sponges had a different aragonite-calcite ratio in comparison to the beach sand sample. However, no species preferentially incorporated particles of particular material type. A wide range of different particles were present in low ratios on the filters (< 3%), such as feldspar, quartz, carbonate phosphate, red pigments and composites. Moreover, several particles are most certainly of anthropogenic origin, i.e., cotton, titanium dioxide, plastic and blue pigments, at ratios between 0.091 to 0.612 particle/mg dry sponge tissue.

### I. Incorporation of foreign particles

The capture and retention of foreign particles are common practice amongst sponges, especially noticeable within members of the subclass Keratosa, for instance species of the genus *Dysidea* embed particles in their spongin fibers (Willenz and van de Vyver 1982; Teragawa 1986a; Cerrano et al. 2007). The pathway most likely used to incorporate coarse particles in the core of spongin fibers is the endocytosis by exopinacocytes (Willenz and van de Vyver 1982; Teragawa 1986a). The diffusion of foreign particles through the ectosome towards the mesohyl in keratose specimens from Bangka Island suggests a particle transfer from the superficial region of the sponge towards the inner one. These findings indicate that the mesohyl serves as a transit zone for particle transport in keratose demosponges. Similar pathways are likely used by heteroscleromorph demosponges; however, patterns observed in our study show differences between accumulation areas. Indeed, Tethyid clades present a thick and dense ectosome composed of organic and inorganic particles of size equal to or smaller than 50 µm diameter. Similar particles also aggregate around choanocyte chambers, which indicates that particles are incorporated via both processes, i.e., captured by the exopinacocytes and absorbed or phagocytized by choanocytes. Consequently, sponges tend to select more voluminous particles (> 50 µm) to support their skeleton in keratose demosponges and to retain smaller particles (< 50 µm) in the ectosome in spiculated demosponges (Teragawa 1986b; Cerrano et al. 2007). Based on these findings, microparticulate pollutants are incorporated by sponges either in skeletal fibers or the ectosome, or both depending on the particle size.

### II. Origins of inorganic particles

#### Scarce minerals

Geologically speaking, North Sulawesi (Indonesia) is a “Neogene island arc […] built upon volcanic-sedimentary basement and underlain by oceanic crust” (Kavalieris et al. 1992); in other words, geological formations are mainly composed of basaltic and andesitic rocks including minerals such as feldspar (Carlile et al. 1990). Intrusion of granite and granodiorite coupled with small localized quartz diorite stocks in the basement of North Sulawesi mainland (Kavalieris et al. 1992) might explain the abundance of fine grained quartz particles reported in all analyzed sponges. Moreover, Manado city and further North, i.e., Bunaken and Bangka islands, is formed with andesitic epiclasts, reworked basaltic rocks rich in quartz and feldspar minerals (Kavalieris et al. 1992). Slightly further North of Bangka island, a modern ferric formation was described by Heikoop et al. (1996) near Mahengetang island (Indonesia). The latter, coupled with commonly reported iron-associated precipitations in shallow-marine environments (Taylor and Macquaker 2011), provide an explanation of the presence of different iron minerals in our samples. Ions of trace metals, such as Pb^2+^, were measured on the coastal area of East Java Province (Indonesia), which might explain the presence of mineral cotunnite encountered in our samples (Apriani et al. 2018). Although a wide range of different mineral compounds have been identified in our study, e.g., magnetite, ferrosilite, cotunnite and mackinawite, the sediment composition on, for example, the Spermonde shelf (South Sulawesi, Indonesia) revealed only hard parts of calcifying organisms (aragonite and calcite) of sizes greater than 250 µm (Janßen et al. 2017). The latter findings however correlate with the abundance of calcium carbonate found in our sand and sponge samples. Although not significantly different from the aragonite-calcite ratio of the beach sand, all sponges incorporated on average more aragonite. We speculate that the surrounding fine suspended matter in the reef might be therefore slightly different than the matter sedimenting in the intertidal range. Nonetheless, our study unravels a wide variety of very fine inorganic particles (< 200 µm) present in the immediate benthic environment, but absent in the sediment (or undetectable with traditional sediment sampling method).

#### Red pigments

Multiple red, orange and pink colored particles that are referred to as “red biomineralization” were present in the beach sand sample and in the 15 specimens analyzed for their particle content. Red pigments of three types (I, II and III) were observed with Raman spectroscopy (Fig. 3). Type I highly resembles the signature of red sclerites from *Tubipora musica. Tubipora* species were indeed an important part of the octocoral fauna at Coral Eye house reef (EB Girard, personal observation). Bergamonti et al. (2011) studied different Raman spectra of various hard-skeleton animals, including *Tubipora musica* and *Corallium rubrum*, and obtained the same vibrational bands as the red pigment type I. The hydrocoral *Stylaster* sp. also exhibits a red pigment, which is identical to that of type II measured on some filters (Bergamonti et al. 2011). Bergamonti et al. (2011) and Thompson et al. (2015) advance that this pigment is linked to carotenoids. The red pigment type I comes from non-methylated polyenes, whereas type II is likely the result of methylated polyenes (Bergamonti et al. 2011; Thompson et al. 2015). However, the origin of these pigments is yet not fully understood. The spectrum of the red pigment type III resembles that of the shell of the bay scallop *Argopecten irradians* (Thompson et al. 2014, 2015). However, this bivalve species is mainly known from the East coast of the United States. Other coral reef organisms from Indonesia most probably have an analogous Raman spectrum, which explains its identification in a few sampled sponges.

### III. Origins of micropollutants

A wide range of different materials was observed in sponges from Bangka Island. On the one hand, they are autochthonous and reflect geological formations, e.g., quartz and feldspar (Carlile et al. 1990; Kavalieris et al. 1992), and reef assemblages, e.g., tunicates and reef-building corals (Yamano et al. 2000; Bergamonti et al. 2011; Łukowiak 2012), of the surroundings. Indeed, slow weathering and erosion processes generate the detachment of particles that compose, together with the reef’s coral sand production, most of today’s Bangka Island sand. On the other hand, they are allochthonous and foreign to the natural environment, such as titanium dioxide, blue-pigmented particles and microplastics.

#### Titanium dioxide

Titanium dioxide (TiO_2_) forms in three minerals: anatase, brookite and rutile, which have differing crystallographic organization (Balachandran and Eror 1982). This aspect enables the identification of the oxide with Raman spectroscopy because different crystalline structures result in distinct Raman signatures of their lattice modes (Balachandran and Eror 1982). Titanium oxides can naturally accumulate in the sand subsequent to weathering of the titanium-bearing mineral ilmenite by underground water (Premaratne and Rowson 2003). However, anatase is also used as a white pigment, i.e., PW6 (titanium white), in automotive paints (Zięba-Palus and Michalska 2014), pharmaceutical coatings (Alexander 2008), thermoplastic resin (Kitamura and Mitsuuchi 1996) and archeological paints (Middleton et al. 2005). Nanoparticulate anatase, together with rutile, is also extensively used for its chemical properties and UVB protective behavior in sunscreens (Yue et al. 1997; Jaroenworaluck et al. 2006; Serpone et al. 2007). Hence, anatase particles found in sponges from Coral Eye house reef might as well come from the degradation of anthropogenic anatase-containing products, and not only from natural sources. The effect of nanoparticulate anatase on health was investigated in mice and corals since it infiltrated the natural environment and its ecotoxicological impact, although not fully understood, concluded little to no effects on health at low concentrations (Duan et al. 2010; Adler and DeLeo 2020).

#### Blue pigments

Particles of highly similar blue color to the human eye were incorporated in two specimens of the clade Tethyid I. The blue pigments, however, showed two different Raman signatures. The main vibrational bands were also measured by Zieba-Palus and Michalska (2014); the authors identified Raman vibrational bands of blue pigments used in car paints. The blue pigment Type I is most probably a mix between the pigment PV23 (dioxazine violet) and PB15 (phthalocyanine 15) and the blue pigment Type II PB15, according to the findings of Zieba-Palus and Michalska (2014). These synthetic organic pigments might also be used in marine coating or recreational painting (Bouchard et al. 2009). Because the Raman vibrational bands of the pigments overwrite that of its polymer composition, it is not possible to identify the material of the particle. However, the pigment PB15 was previously recorded as a dye associated with microplastics isolated from the soft tissue of bivalves (van Cauwenberghe and Janssen 2014) and intertidal textile fibers (Girard et al. 2020).

#### Textiles and microplastics

High tides and winds bring large quantities of marine debris, including plastics and textiles, on Bangka Island (EB Girard, personal observation; (Giebel 2018) unpublished report). The litter lands on beaches, where the highest degradation rate of plastic has been reported (Andrady 2017). Coral Eye Resort volunteers clean the beach daily, however this is not done systematically all around the island yet, nor on proximal coast lines (EB Girard, personal observation). Consequently, eight microparticles were herein identified as cotton (n = 6) or polystyrene (n = 2) in all sampled keratose species. Cotton has also been reported to be the most observed fabric in environmental dust, as fibers in the atmosphere, but also in the intertidal zone (Dris et al. 2016, 2017; Girard et al. 2020). Nevertheless, some of these particles may also originate from the cloth made of cotton that was used to dry glass dishes to avoid plastic contamination of the samples. Polystyrene is one of the three most abundant microplastic materials reported at sea, together with polyethylene and polypropylene (Andrady 2017; Auta et al. 2017). Our results are further supported by the findings of Ling et al. (2017), who estimated that the concentration of microplastic particles in the sediment reaches up to 0.4 particle/mL in the southern coasts of Australia. The authors noticed a consistent microplastic concentration across 42 sampling sites. Moreover, microparticles of plastic were at highest concentration in a size range ca. 60-400 µm (Ling et al. 2017), which is concordant with the particle size incorporated by the sponge exopinacoderm. Because sponges can pump several decades to hundreds of liters per day (Leys et al. 2011) and microparticles deposit on their ectosome, Bangka specimens indeed incorporated microplastics.

### IV. Sponges as bioindicators

Sponges are potentially ideal local bioindicators because they are sessile animals and widely distributed across all aquatic habitats. In fact, the families of sponges occurring around Bangka island have previously been recorded from various other localities in Indonesia (van Soest 1989, 1990; Cerrano et al. 2002, 2006; Bell and Smith 2004; de Voogd et al. 2006, 2009; de Voogd and Cleary 2008; Becking et al. 2013; Calcinai et al. 2017a, b) (Supplementary material Fig. S4). Furthermore, a recent study identified sponges as a good monitor to record more efficiently DNA of surrounding vertebrates than robotic samplers for environmental DNA (eDNA) (Mariani et al. 2019). Sponges have also been recognized as bioindicators for environmental stress (Carballo et al. 1996), water quality (Mahaut et al. 2013), and multiple pollutants, e.g., heavy metal pollution (Selvin et al. 2009; Venkateswara Rao et al. 2009) and polychlorobiphenyl (Perez et al. 2003). Collection sponges have also been recently surveyed for fibrous microplastics (Modica et al. 2020).

#### Extrapolation to realistic micropollutant concentrations

Based on our results, sampled sponges did not preferentially incorporate particles of specific materials, which suggests that fluctuation in material ratios is due to the spatial variation of surrounding microparticles. At a concentration higher than 0.1 particle/mg of dry sponge tissue, here from keratose demosponges (*Carteriospongia* sp. and *Icrinia* spp.) weighing ca. 6 mg (dry weight), we extrapolate that at least 10,000 microplastic particles can be incorporated by sponges weighing more than 100 g (dry weight). Similar approximations can derive from the results regarding abundance of blue-pigmented particles, cotton and titanium dioxides in the spiculated demosponge Tethyid species. Because sponges can weigh several hundreds of grams (Reiswig 1971; McMurray et al. 2008), they have the potential to accumulate non-negligible amounts of micropollutants. A larger screening of associated particles in sponge tissue is likely to reveal more microplastics and other particles derived from anthropogenic products.

Sponges are therefore efficient sediment traps, recording the diversity of the matter in the ambient water as they are able to register this diversity to the finest grain (< 200 µm), otherwise difficult to recall solely based on sand samples (Janßen et al. 2017), as observed in this study. Such biological monitors also provide information over a time window, whereas traditional net tows represent only a single point in time (Lindeque et al. 2020). Furthermore, in sediment traps, the fine fraction is washed away by currents, leading to biases in the actual particle diversity present in the sediment at a given location (Janßen et al. 2017). Based on current knowledge and results from this study, we conclude that sponges also have the potential to biomonitor micropollutants, such as particles putatively originating from anthropogenic products (e.g., microplastics).

### VI. Outlook

As sponges incorporate micropollutants from their surroundings, a sample of sponge tissue may provide a unique estimate of the local micro-pollution applicable to the immediate fauna. This study also narrows the knowledge gap on particle incorporation processes and provides a first assessment on the particle diversity in sponges. For future research on the topic, we suggest analyzing a large amount of tissue and particles by, for instance, dissolving the carbonates, which will provide a higher resolution of the diversity of scarce particles incorporated by sponges.

Further hypothetical outcomes arise as sponges incorporate micropollutants, for instance toxins associated with these microparticles can leach, impacting sponge development and pumping capacity (Hill et al. 2002). Likewise, microbial pathogens hitchhiking on, for example, microplastics may negatively affect sponges (Taylor et al. 2007), both of which will have a direct impact on the ecosystem they inhabit. Keratose demosponges may also use particulate micropollutants to build their skeleton and support their growth, creating temporary sinks or an expressway to enter the marine food chain through spongivores. On a more positive note, sponges likely host degrading bacteria able to remineralize certain micropollutants (Lee et al. 2001), taking the sponge loop theory to the next level.

## ACKNOWLEDGEMENTS

We thank the Indonesian authorities for providing the research visa and permit (research permit holder: Elsa Girard; SIP no.: 97/E5/E5.4/SIP/2019) to conduct the research activities on Bangka Island, in collaboration with Sam Ratulangi University Manado (UNSRAT, Indonesia). We thank Dirk Erpenbeck, Oliver Voigt, Anna Clerici, Marco Perin, Stefanie Ries, Magdalena Wilde and Samuel Leivy Opa for helping during field work activities. A last word to thank the No-Trash Triangle Initiative for tackling one of the big problems Earth is facing today.

