## Supplementary material for "Sponges as bioindicators for microparticulate pollutants"

### I. Supplementary figures

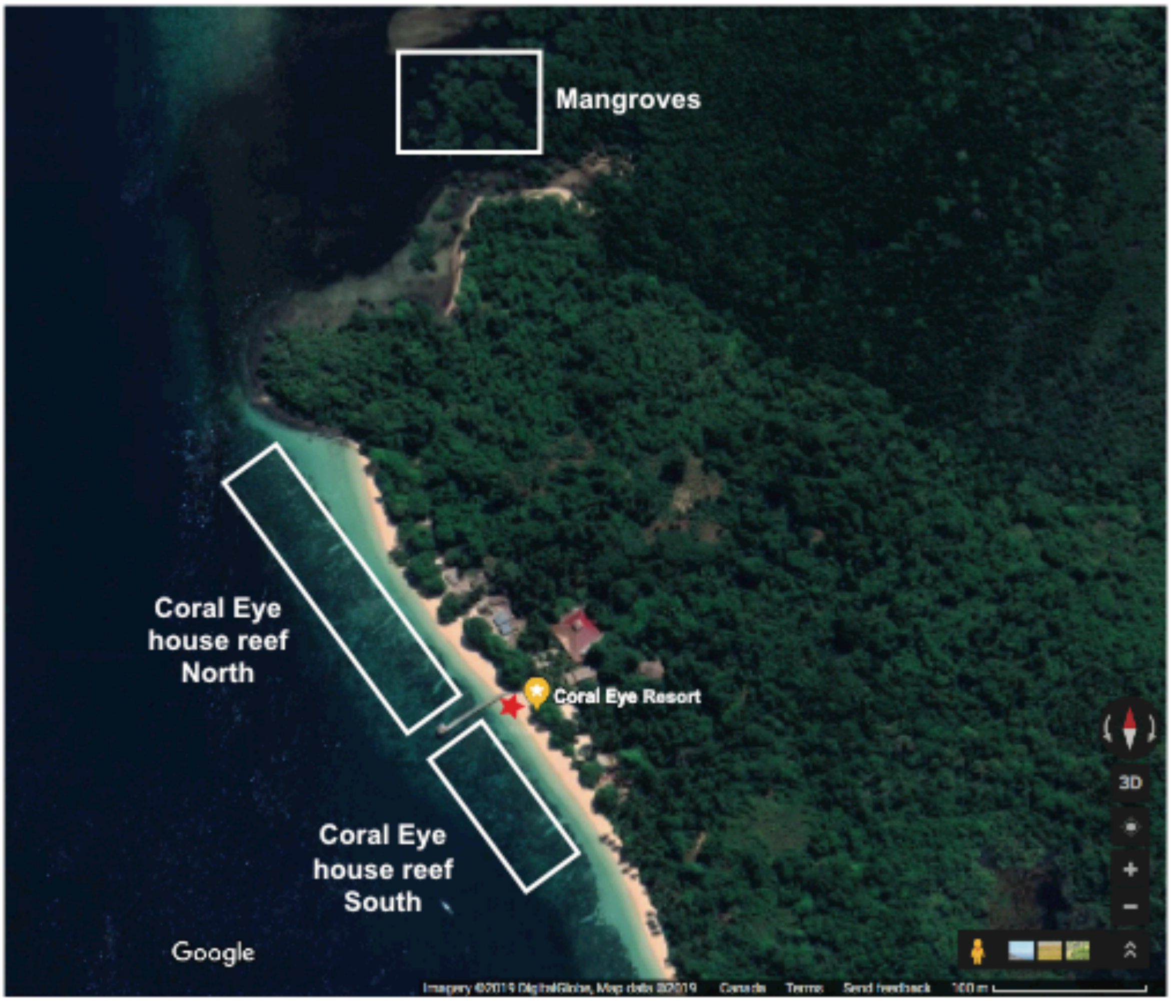


**Figure S1.** Detailed sampling site on the west coast of Bangka Island (North Sulawesi, Indonesia). Specimens were sampled along Coral Eye house reef and at the nearby mangroves. The location of the beach sand sample is marked with a red star.

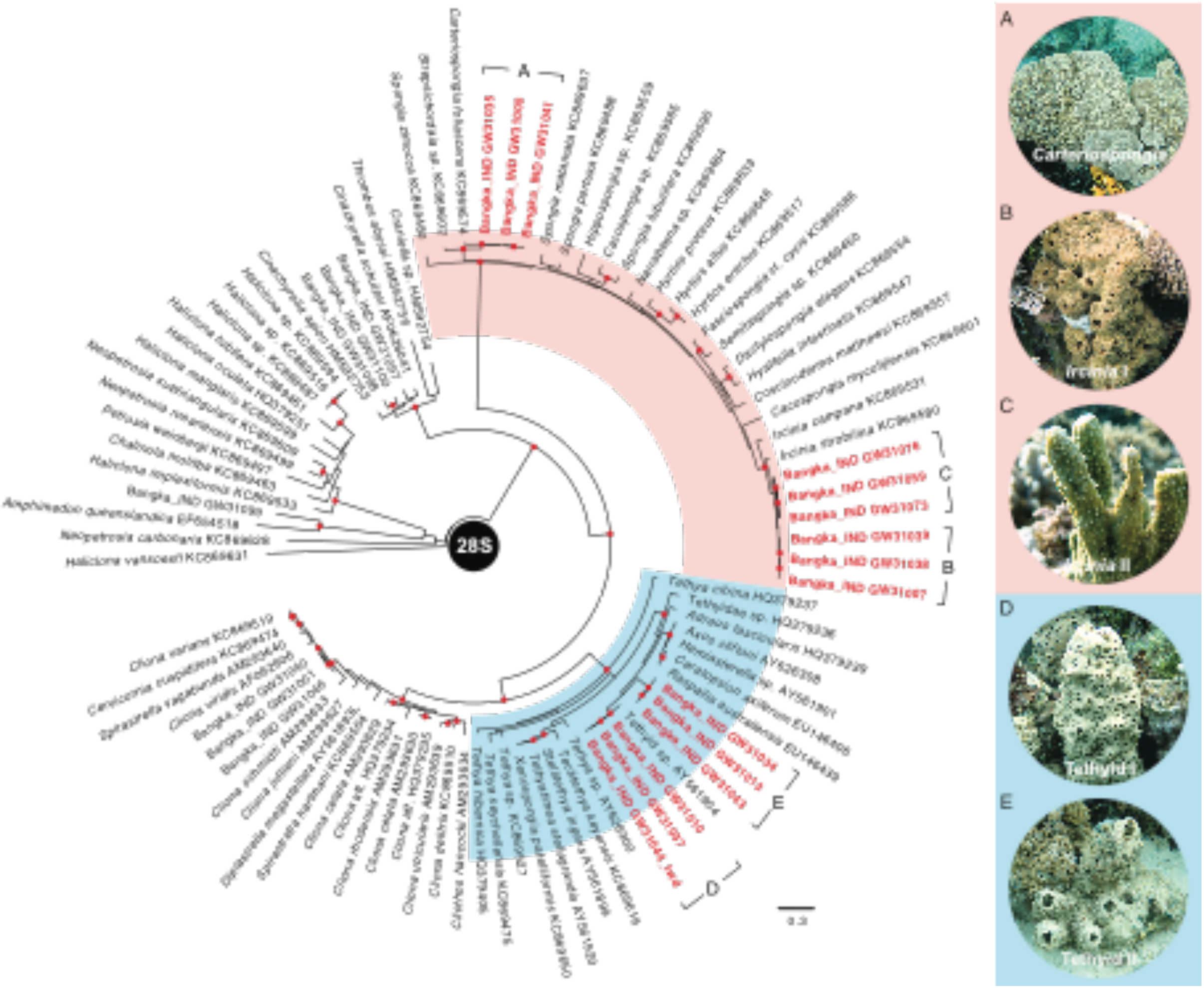


**Figure S2.** Phylogenetic reconstruction of successfully DNA barcoded specimens. A) phylogenetic reconstruction of 28S sequences. The orange clade highlights all samples of the subclass Keratosa (class Demospongiae), while the remaining species are part of the subclass Heteroscleromorpha (class Demospongiae). The square parentheses (A, B, C, D, E) indicate to which clade it is referred to. Pictures illustrate the marco-morphology associated with each triplicate (*Carteriospongia* (A); *Ircinia* I (B) and II (C); Tethyid I (D) and II (E)). Samples collected and analyzed in this study are highlighted in red. Bootstrap > 70 is marked with a red dot.

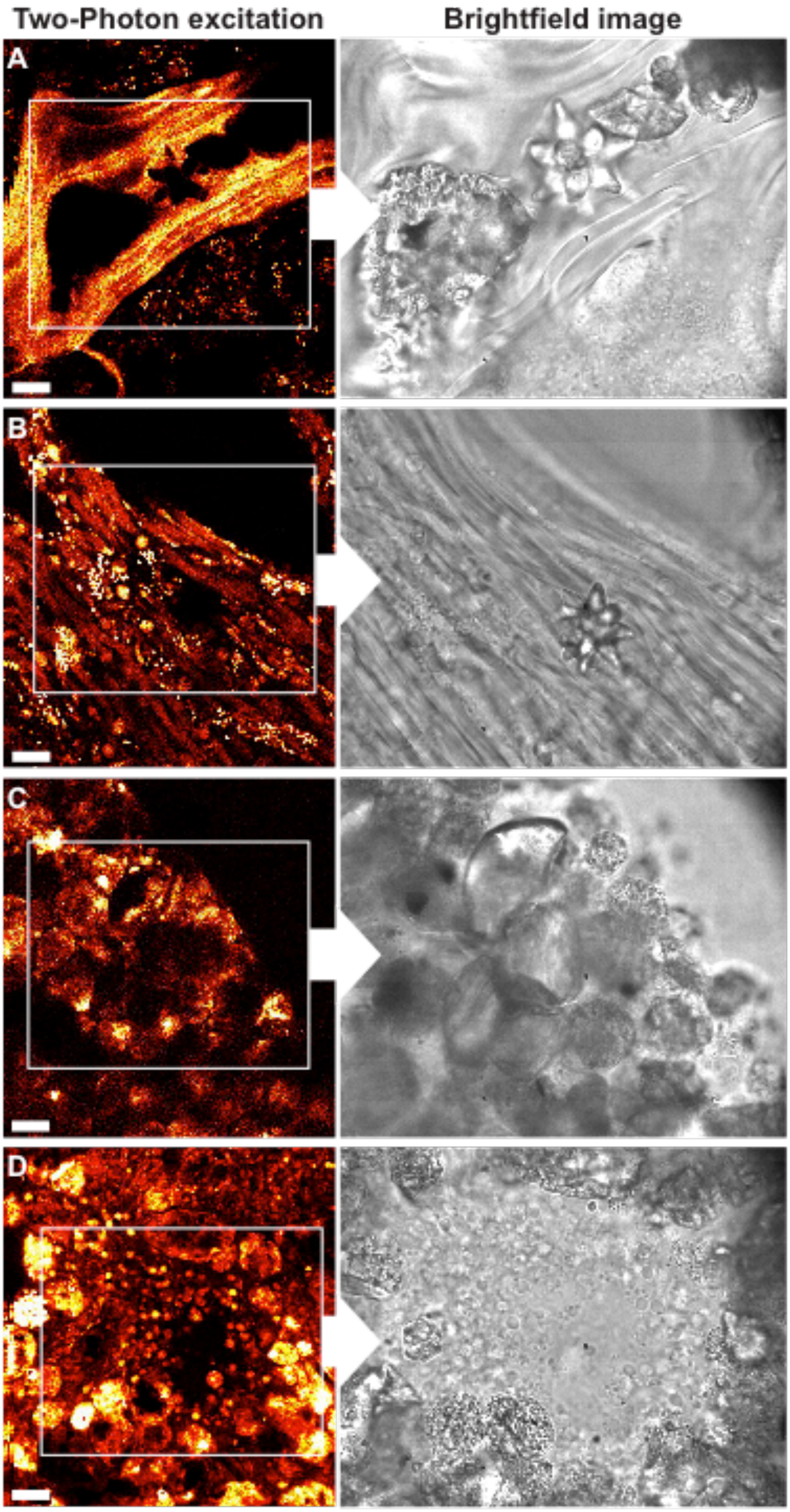


**Figure S3.** Two-photon excitation images combined with their associated brightfield images. Exogenous particles embedded in the tissue are found A) in spongin fibers of *Carteriospongia* sp., B) in mesohyl of *Ircinia* sp., C) at the ectosome of Tethyid sp., and D) surrounding the choanocyte chamber of Tethyid sp.. Two-photon excitation images are average intensities over the z-range. Scale bars are 20 µm.

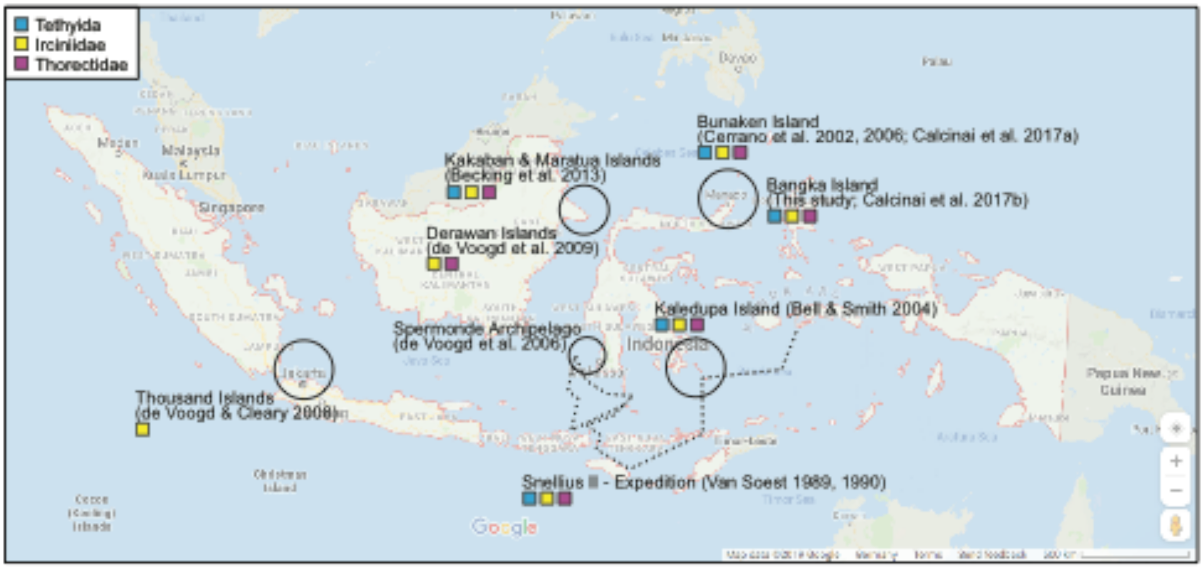


**Figure S4.** Summary of the distribution of identified families at Coral Eye house reef across Indonesia. Note: the record of marine sponge biodiversity in Indonesia is limited to some publications. Indonesia is outlined in pale red.

### II. Supplementary tables

**Table S1.** Details regarding the sample collection at Coral Eye Resort, North Sulawesi, Indonesia (sample identification number, sampling date and location).

| Sample ID | Sampling date | Sampling location | Jetty side |
|---|---|---|---|
| GW31000 | 17.03.2019 | Coral Eye Housereef | South |
| GW31001 | 17.03.2019 | Coral Eye Housereef | South |
| GW31007 | 30.03.2019 | Coral Eye Housereef | South |
| GW31010 | 30.03.2019 | Coral Eye Housereef | South |
| GW31013 | 01.04.2019 | Coral Eye Housereef | South |
| GW31019 | 03.04.2019 | Coral Eye Housereef | South |
| GW31022 | 03.04.2019 | Coral Eye Housereef | South |

| | | | |
|---|---|---|---|
| GW31034 | 05.04.2019 | Coral Eye Housereef | South |
| GW31035 | 05.04.2019 | Coral Eye Housereef | South |
| GW31037 | 05.04.2019 | Coral Eye Housereef | South |
| GW31038 | 05.04.2019 | Coral Eye Housereef | South |
| GW31039 | 05.04.2019 | Coral Eye Housereef | South |
| GW31040 | 05.04.2019 | Coral Eye Housereef | South |
| GW31041 | 05.04.2019 | Coral Eye Housereef | South |
| GW31042 | 05.04.2019 | Coral Eye Housereef | South |
| GW31043 | 05.04.2019 | Coral Eye Housereef | South |
| GW31044 | 05.04.2019 | Coral Eye Housereef | South |
| GW31045 | 05.04.2019 | Coral Eye Housereef | South |
| GW31046 | 05.04.2019 | Coral Eye Housereef | South |
| GW31058 | 06.04.2019 | Coral Eye Housereef | North |
| GW31059 | 06.04.2019 | Coral Eye Housereef | North |
| GW31071 | 07.04.2019 | Coral Eye Housereef | North |
| GW31073 | 07.04.2019 | Coral Eye Housereef | North |
| GW31076 | 07.04.2019 | Coral Eye Housereef | North |
| GW31077 | 09.04.2019 | Mangroves | --- |
| GW31079 | 09.04.2019 | Mangroves | --- |
| GW31084 | 09.04.2019 | Mangroves | --- |
| GW31085 | 09.04.2019 | Mangroves | --- |
| GW31096 | 09.04.2019 | Mangroves | --- |
| GW31097 | 09.04.2019 | Mangroves | --- |
| GW31098 | 09.04.2019 | Mangroves | --- |
| GW31099 | 09.04.2019 | Mangroves | --- |
| GW31100 | 09.04.2019 | Mangroves | --- |

**Table S2.** Primer list used to amplify 28S, their associated PCR regime and reagent details (forward (fwd); reverse (rev)).

| **Reagent recipe (1 reaction)** | | MilliQ 8.9 µL, PCR Flexi Buffer 5x 5 µL, $MgCl_2$ (25 mM) 3 µL, dNTP (10 mM) 1 µL, Forward primer 1 µL, Reverse primer 1 µL, Taq polymerase 0.1 µL and DNA (2-10 ng/µL) 2 µL | | |
|---|---|---|---|---|
| **Primer #** | **Primer name** | **Target gene** | **Primer sequence (5'-3')** | **References** |
| 12 | 28S-C2-fwd | 28S rDNA fwd | GAAAAGAACTTTGRARAGAGAGT | Chombard et al. 1998 |
| 13 | 28S-D2-rev | 28S rDNA rev | TCCGTGTTTCAAGACGGG | |
| **Gene** | **PCR program** | | | |
| 28S | Denaturation at 96 °C (3 min) → 35 amplification cycles: denaturation at 96 °C (30 s), annealing 50 °C (30 s), elongation 72 °C (30 s) → elongation at 72 °C (5 min) | | | |

**Table S3.** Triplicates and their associated classification based on barcoding. Subclass Keratosa (A, B and C), Subclass Heteroscleromorpha (D and E). Order and Family were taken from the World Register of Marine Species (WoRMS - http://www. marinespecies.org/).

| Triplicate | Associated samples | Clade name | Classification (Phylum Porifera, Class Demospongiae): Order - Family - Genus |
|---|---|---|---|
| A | GW31000-35-41 | *Carteriospongia* | Dictyoceratida - Thorectidae - *Carteriospongia* |
| B | GW31007-38-39 | *Ircinia* I | Dictyoceratida - Irciniidae - *Ircinia* |
| C | GW31059-73-76 | *Ircinia* II | Dictyoceratida - Irciniidae - *Ircinia* |
| D | GW31010-37-44 | Tethyid I | Tethyida - unclassified Tethyida |
| E | GW31013-34-43 | Tethyid II | Tethyida - unclassified Tethyida (cf. *Liosina*) |

**Table S4.** List of different particles, which could be unequivocally identified, associated with their vibrational bands. Vibrational band values in bold indicate the dominant intensity peaks characterizing the material signature. "NA" stands for "not applicable".

| **Material** | **Formula material** | **Formula pigment** | **Vibrational bands (0-2000 $cm^{-1}$)** | **Reference for identification** |
|---|---|---|---|---|
| **Aragonite** | $CaCO_3$ | NA | **151**, 202, 701, **1083** | RRUFF (ID R060195; R080142) |
| **Calcite** | $CaCO_3$ | NA | 155, **281**, 712, **1088** | RRUFF (ID R040070) |
| **Quartz** | $SiO_2$ | NA | 126, 200, 262, 355, **461** | RRUFF (ID R050125) |

| | | | | |
|---|---|---|---|---|
| **Anatase** | $TiO_2$ | NA | **141**, 194, 390, 510, 630 | RRUFF (ID R070582) |
| **Feldspar (Anorthite)** | $Ca(Al_2Si_2O_8)$ | NA | 162, **290**, 411, **480**, **509** | RRUFF (ID R050104) |
| **Carbonate phosphate** | $CO_7P_5$ | NA | **960**, 1073 | Awonusi et al. 2007; Slimani et al. 2017 |
| **Carbon** | a-C | NA | 1349, 1593 | Milani et al. 2015 |
| **Graphite** | C | NA | 1357, **1580**, 1621 | Malard et al. 2009; Milani et al. 2015 |
| **Red type I (pigment canthaxanthin)** | $CaCO_3$ | $C_{40}H_{52}O_2$ | 1021, **1126**, 1301, **1511** | Bergamonti et al. 2011 |
| **Red type II** | $CaCO_3$ | Not identified | 974, 1019, 1090, **1161**, 1209, 1453, **1523** | Gaspard at al. 2019 |
| **Cotton** | $(C_6H_{10}O_5)_n$ | Not identified | 329, **377**, 432, 459, 520, 898, 999, **1093**, **1121**, 1152, 1339, **1380**, 1411, 1478 | Rygula et al. 2011; Grishanov 2011 |
| **Blue type I (pigment PV23 dioxazine violet)** | Not identified | $C_{34}H_{22}Cl_2N_4O_2$ | 320, 489, 531, 592, 621, 683, 921, 1149, 1169, **1210**, 1258, **1347**, **1392**, **1431**, **1530**, 1597 | Ziebas-palus et al. 2014 |
| **Blue type II (pigment PB15 Copper phthalocyanine)** | Not identified | $C_{32}H_{16}CuN_8$ | 591, 680, 1084, 1142, **1340**, **1450**, **1528** | Ziebas-palus et al. 2014 |
| **Polystyrene** | $(C_8H_8)n$ | NA | 621, 810, **1001**, 1032, 1089, 1156, **1450**, 1605, 1666, 1730 | Palm 1951; Mazilu et al. 2007 |
| **Red type III (shell pigment)** | $CaCO_3$ | Not identified | 1118, 1502 | Thompson et al. 2015 |
| **Magnetite** | $Fe_3O_4$ | NA | 157, **199**, **212**, 290, 510, **660**, 1310 | de Faria et al. 1997; Hanesch 2009 |
| **β-Quartz** | $SiO_2$ | NA | 112, **458** | McMillan et al. 1992 |
| **α-quartz with fulgurite** | | | | |
| **Shocked α-quartz** | | | | |
| **Mackinawite** | FeS | NA | **217**, 283 | Boughriet et al. 1997; Bourdoiseau et al. 2008; Genchev and Erbe 2016 |
| **Ferrosilite** | $Fe^{2+}{}_2Si_2O_6$ | NA | 305, 379, 655, 993 | RRUFF (ID R070386; X050187); Huang et al. 2000; Wang et al. 2001 |
| **Hematite** | $\alpha$-$Fe_2O_3$ | NA | 210, 239, 290, **406**, 600 | de Faria et al. 1997; Jubb and Allen 2010 |
| **Riebeckite-derivative** | $Na_2(Fe^{2+}{}_3Fe^{3+}{}_2)Si_8O_{22}(OH)_2$ | NA | **92**, 242, 350, **448**, 505, 562, **599**, 881, 910, 1083 | Apopei et al. 2011 |
| **(Amorphous) Calcite** | $CaCO_3$ | NA | 337, 369, 1086 | Porto et al. 1966; Dawson et al. 1973; De |

| | | | | |
|---|---|---|---|---|
| **Calcium hydroxide** | $Ca(OH)_2$ | | | La Pierre et al. 2014; Schmid & Dariz 2015 |
| **Cotunnite** | $PbCl_2$ | NA | 157 | RRUFF (ID R060655) |
